# Latching dynamics as a basis for short-term recall

**DOI:** 10.1101/2021.02.18.431782

**Authors:** Kwang Il Ryom, Vezha Boboeva, Oleksandra Soldatkina, Alessandro Treves

## Abstract

We discuss simple models for the transient storage in short-term memory of cortical patterns of activity, all based on the notion that their recall exploits the natural tendency of the cortex to hop from state to state – latching dynamics. We show that in one such model, and in simple spatial memory tasks we have given to human subjects, short-term memory can be limited to similar low capacity by interference effects, in tasks terminated by errors, and can exhibit similar sublinear scaling, when errors are overlooked. The same mechanism can drive serial recall if combined with weak order-encoding plasticity. Finally, even when storing randomly correlated patterns of activity the network demonstrates correlation-driven latching waves, which are reflected at the outer extremes of pattern space.

## 1 Introduction

If short-term memory (STM) is expressed by the activity of the same neurons that participate in the representation of long-term memories (LTM) [1], what determines its drastically reduced capacity? What makes us able to recognize tens of thousands of images as familiar [2] and yet unable to detect a change in a configuration of more than a few elements that we have just seen [3]? Focusing on the recall of sequences of well-known items, what makes it so difficult to go, again, beyond very short sequences?

Within the general hypothesis that interference between memories is critical [4], we consider the specific hypothesis that the imprecise mechanism for short-term storage and the necessity to rely on intrinsic, *mind-wandering* neural dynamics for retrieval and recall [5] can make either interference from items in long-term memory or the randomness in retrieval trajectories the limiting factor, depending on the task.

Taking the cue from a statistical theory of trajectories among ensembles of items [6], we analyse latching dynamics expressed by a Potts network, taken as a model of the interactions among patches of cortex, when it hops spontaneously from activity pattern to activity pattern, recalling them in a sequence resembling a random walk.

We show that by adding a mechanism that gives an extra “kick” to a small subset of *L* among *p* patterns in long-term memory, latching dynamics can be approximately restricted to the subset, which are effectively kept in short-term memory. The usefulness of such partially spontaneous dynamics is critically limited by the nature of the short-term task it sub-serves, in agreement with established results and with our own observations in tasks requiring short-term memory of spatial locations. In free recall, where repetitions and mistakes are not penalised, the number *M* of retrieved items tends to scale sublinearly with *L*, reflecting largely random exploration. In a task which is terminated by mistakes, instead, capacity is constrained by the interference of other items in long-term memory, with a limit dependent on the type of short-term kick. In contrast, modeling serial recall with hetero-associative short-term synaptic enhancement leads to guided dynamics, which if the enhancement is arbitrarily effective can outperform human subjects, but also abolish latching dynamics. We discuss what this may imply for actual serial recall mechanism. While the Potts network does not have the articulation that would allow fully benchmarking its validity as a model of short-term memory [7], we believe that it can help clarify some of the critical factors that limit its effectiveness.

The paper is organized as follows. In section 2, we compare three models for short-term memory. In section 3 we look at the performance of our second model for free recall and compare it with experimental data. Modeling serial recall is discussed in section 4 and the quasi-random walk trajectories followed by latching sequences are discussed in section 5.

## 2 Simple mechanistic models for short-term memory

If latching dynamics is effectively constrained to a small subset of the activity patterns that represent items in long-term memory, it can serve a short-term memory function. Different neural-level mechanisms can constrain the dynamics, and it can be envisaged that several of them may operate in synergy. Here we analyse three, which can be simply associated with distinct parameters of the Potts network, and we consider each mechanism separately from the other two, to demonstrate its characteristics (Fig. 1a). The parameters we focus on are the degree of local feedback (Model 1), the local adaptive thresholds (Model 2) and the strength of long range connections (Model 3).

**Figure 1:**
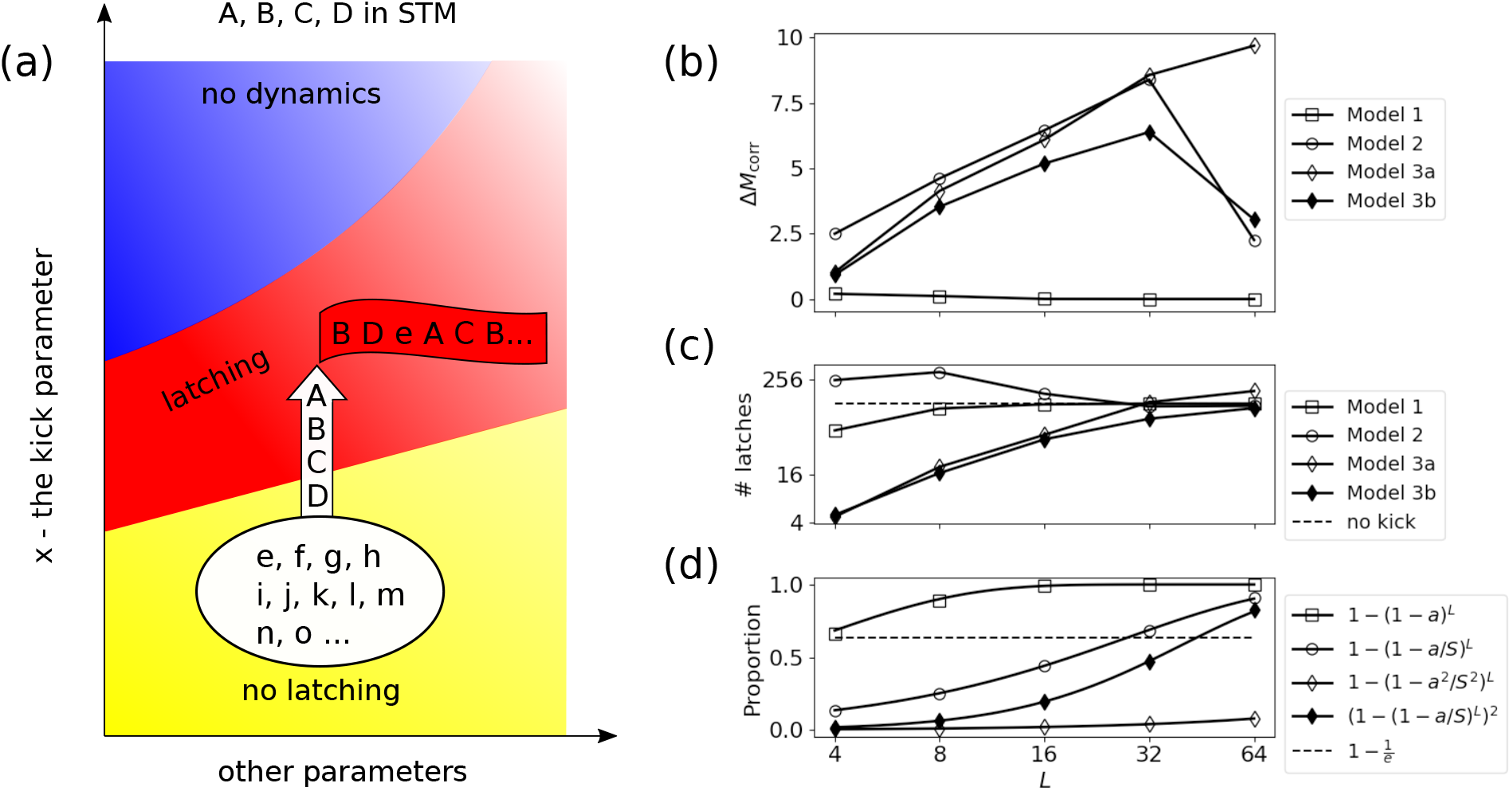
Different models for holding items in STM yield qualitatively different recall performance. **(a)**: Schematic diagram of models for STM. The STM function is produced by a“kick” Δ*x* in the parameter *x*, representing *w*, *θ* and *J* for Model 1, 2, and 3, respectively. **(b)** The quantity Δ*M_corr_* has a maximum at around *L* ≃ 32 for Model 2 and 3b and it continues to grow for Model 3a, while it remains always close to zero for Model 1. The abscissa is *L*, the number of items in STM, in log scale. The ordinate is Δ*M_corr_ M*_corr_(Δ*x* = 0.3) *M*_corr_(Δ*x* = 0.0), where *M*_corr_ is the number of recalled STM items until the network either repeats an already-visited item or (mistakenly) retrieves one of the LTM items. **(c)** The different propensity to latch, i.e., to make transitions, is quantified by the number of latches p er sequence, plotted as a function of *L* for the 3 models, in a log-log scale. The strength of the kick is, again, Δ*x* = 0.3 for each model. The horizontal dashed line indicates the number of latches per sequence when all *p* patterns are on equal footing, i.e., the strength of the kick is 0. **(d)** The proportion of resources utilised in the models predicts the peak of the performance Δ *M_corr_*. The dashed horizontal line indicates the proportion equal to 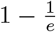. Across all 3 panels, parameters are *p* = 200, *S* = 7, *a* = 0.25, *γ_A_* = 0.5 and *w* = 1.1.

In each case, we change therefore a single parameter across many network elements, so that *L* patterns, those supposed to be held in short-term memory, are driven into the latching regime. This change, which embodies short-term *storage*, should avoid pushing into the latching regime also the other *p* − *L* patterns, but to some extent their involvement is unavoidable, as will be shown.

First, we describe how in principle the Potts model for long-term memory can express also a short-term memory function. In practice, for the sake of a fair comparison among the mechanisms, in the simulations we change each parameter the other way around, as explained in 2.4 below.

### 2.1 Model 1: Stronger local feedback for the items held in STM

The first mechanism models increased depth of the attractors in the patches of cortex where any of the *L* patterns is active, which could reflect a generic short-term potentiation of the synaptic connections among pyramidal cells in those patches, what in the Potts network is summarily represented by the parameter *w* ([8, 9]). In the model, each of the *L* items is active over *aN* Potts units, and their active states are shared with many other items not intended to go into STM. This is the coarseness that leads to limited capacity of memory: if *L* is too large, virtually all of the units are given the kick, all with the same strength, and no distinction between the *L* selected items and the other *p* − *L* remains. Formally, instead of common *w* for all Potts units, we introduce

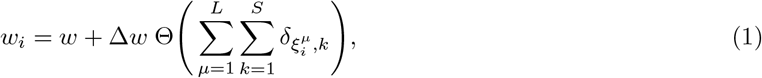

where 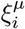 is the state of pattern *ξ^μ^* at the unit *i*, Θ(·) is the Heaviside step function and 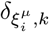 is the Kronecker’s delta symbol.

If a unit participates in the representation of any one of the *L* patterns in STM, then *w_i_* = *w* + Δ*w*. If not, *w_i_* = *w*.

### 2.2 Model 2: Lower adaptive threshold for the items held in STM

In the second mechanism, a parameter regulating firing rate adaptation is reduced selectively for the neurons that are active, in those patches, in the representation of the *L* items. That is, we *decrease* adaptation, by subtracting from the adapted threshold 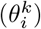 a term Δ*θ*, for the Potts states that are active in any one of the *L* patterns,

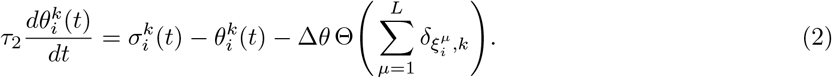

### 2.3 Model 3: Stronger long-range connections for the items held in STM

The third mechanism we consider is the one acting on the long-distance synaptic connections between neurons, represented in the Potts model [8] by the tensor connections between Potts units. We model short-term potentiation of the synaptic connections by stronger tensor connections. Since the latter connect separate Potts units, however, in order to specify exactly which tensor elements are considered to be potentiated, we have to specify whether the *L* patterns are taken to be stored simultaneously. We consider two opposite cases. If they are assumed to be all stored at separate times, the stronger tensor elements are those that connect Potts states of two units both active in any one of the *L* patterns. If they are assumed to be all stored in STM together, the stronger elements are all those that connect Potts states of two units both active in any *pair* of the *L* patterns. We call them variants *a* and *b* of Model 3.

#### Model 3a: Model 3 with only autoassociative connections in short-term memory

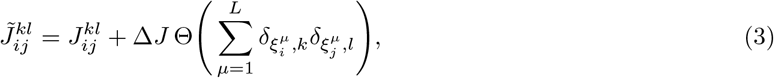

where 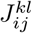 is the strength of connections that do not belong to any one of *L* patterns in STM, given in Eq. (6). Here we say that a connection belongs to a pattern when the two states that are paired by the connection participate in the representation of the pattern.

#### Model 3b: Model 3 with all associative connections among STM items

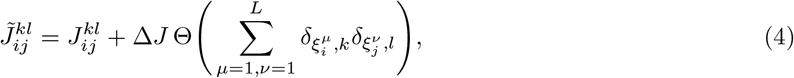

where 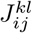 is again given in Eq. (6). In this model, we potentiate extra connections in addition to those that are potentiated in Model 3a. These are the so-called heteroassociative connections that connect Potts states of one item to those of another item in STM.

### 2.4 Comparing the models

For the sake of a fair comparison among the mechanisms (Models 1, 2 and 3), we equalize the values of all parameters as they affect the *L* patterns, so that in practice, rather than bringing them into the latching regime, which is what should happen in the real process, in our model evaluation we push the other *p* − *L* out, or partially out, in different directions.

We first consider how effective are the three mechanisms in constraining latching dynamics to the *L* items in STM. To this end, we define *M* _corr_ as the number of recalled STM items until the network either repeats an already-visited item or retrieves one of the LTM items. We then consider the difference between this quantity and the value it would have without any differentiation between the *L* and the other items, Δ*M*_corr_ ≡ *M*_corr_(Δ*x* = 0.3) − *M*_corr_(Δ*x* = 0.0); this subtraction of the chance level quantifies the genuine effect of Δ*x*. Here *x* represents *w*, *θ* and *J* for the three models, respectively.

We find that for some of the mechanisms, latching dynamics are effectively constrained to the *L* items, but only up to a given value of *L* (Fig. 1b). We can understand this limitation as being due to interference from the LTM items.

Let us consider the proportion of elements (units, states and connections for Model 1, 2 and 3, respectively) that are kicked for a given number *L*. If we randomly pick one unit (state or connection), then the probability of it belonging to one of *L* patterns in STM can be written as *P_L_* = 1 − (1 − *a*)^*L*^ for Model 1. Similarly we have 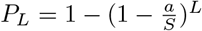 for Model 2, 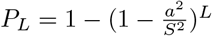 for Model 3a and finally 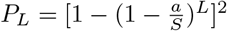 for Model 3b (Fig. 1d). All of these quantities approach 1 when *L* becomes very large. As a rough estimation, we can set a criterion of 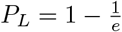, above which one cannot discriminate STMs from LTMs. We can then compute the critical value of *L*, *L_c_*, at which *P_L_* reaches this criterion (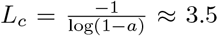 for Model 1, 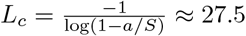 for Model 2, 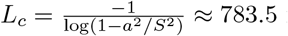 for Model 3a and 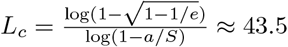 for Model 3b given the parameters for which we run the simulations, *S* = 7, *a* = 0.25).

When we increase *L*, there are two factors that affect *M*_corr_. The first one is that the seemingly quasi-random trajectory among the *L* items lasts longer, which makes *M*_corr_ larger. If this exploration has the characteristics of a random walk, *M*_corr_ should grow like 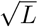 (see Appendix), provided there are no *errors*, i.e. recall of items that are not in STM. The occurrence of errors is the second factor that affects *M*_corr_, causing it to drop as *L* increases beyond approximately the value *L_c_* above. When *L* is small, the first factor dominates and *M*_corr_ grows. Beyond *L_c_*, there is an avalanche of errors as there are many LTM patterns that are kicked as strongly as those in STM. Then the second factor dominates, decreasing *M*_corr_. This semi-analytic consideration thus allows us to predict the positions of the maximum values of *M*_corr_ (Fig. 1b). Indeed, the predicted values of *L_c_* approximately match the positions of the peaks we obtain through simulations, but we can see them only for Models 2 and 3b because for Models 1 and 3a the estimated values of *L_c_* (and thus the positions of the peaks) are too small or large compared to the range of *L* we have considered (Fig. 1b). Nevertheless, we find that the parameter values chosen are representative of a broad range, e.g. for Δ*θ* between 0.1 and 0.5 (Figs. S1 – S4).

It is clear that Model 1 has very limited capacity to constrain latching dynamics, in that interference effects occur already for low values of *L*, and the values of Δ*M*_corr_ are modest compared to those obtained with Models 2 and 3.

In contrast, Models 2 and 3 appear broadly equivalent in terms of Δ*M*_corr_, except that the mechanism acting on the long-range connections, in its variant 3a, is not affected by interference until much higher *L* values. This is because in this case, the kick is more selectively affecting a subset of the very many *N* × *C* × *S* × (*S* − 1)/2 tensor connection values in the model.

There is, however, a major difference between Models 2 and 3 if we consider their overall propensity to latch. To measure this propensity, we first cue the network with one of the memorised patterns, after which we count the total number of latches that occur until the dynamics stop on their own (Fig. 1c). We can see that with Model 2, constraining the dynamics to be among the *L* items actually enhances the length of the latching sequences, in the simulations, representing an increased tendency to latch, whereas the opposite is true, at least up to moderate values of *L*, for Model 3 (and incidentally, for Model 1). This can be qualitatively understood by considering the different roles played by the parameters that we change in the three models. For Model 2 and 3, introducing an extra “kick” does effectively restrict latching dynamics to the STM subset, if *L* is not too large, but it affects the overall propensity of the network to latch in different ways. Suppose *L* = 4. There are only 4 patterns that the network can visit. Because of “neuronal” fatigue mimicked by 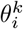, a pattern cannot be revisited soon after it is first visited. Therefore we see a slow increase (with respect to *L*) in latching propensity for Model 3 (Fig. 1c). But for Model 2, the “refractory” effect of 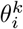 is greatly screened due to the direct manipulation on 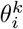. At *L* = 4 for Model 2 we observe clean latching, in the sense that the condensed pattern is almost perfectly retrieved whereas non-condensed patterns have virtually zero overlap (Fig. S5).

In summary, when considering the quantity Δ*M*_corr_ as well as the propensity to latch, Model 2 expresses a functionality more plausibly associated with what we would expect from short-term memory. First, the quantity Δ*M*_corr_ does not increase indefinitely upon increasing *L*, and has a limit. Moreover, the basic propensity to latch also falls off with increasing *L*, reminiscent of the slowing down of retrieval from memory as the set size increases [7]. Therefore, in the remainder of this work, we focus on Model 2.

## 3 Can “free recall” by the Potts network model experimental data?

Having discussed three different models for short-term recall, we study in detail Model 2, that uses a lower adaptive threshold as a mechanism to hold items in STM. We run extensive simulations and see how it functions, comparing it also with experimental data.

### 3.1 If limited by repetitions, the network *can* recall up to 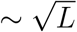 STM items

Experimental data from decades ago show that in some free recall tasks, the number of items recalled from memory obeys a power law of the list length [2, 10]. To explain this finding and more generally to investigate the putative mechanisms that hinder recall, a conceptual model for memory recall has been proposed [6, 11]. In this model, a quantity *R* is defined as the number of recalled items until the searching trajectory enters a loop, which is then iterated indefinitely. STM items are drawn, in their framework, from a virtually unlimited reservoir of (LTM) memory items. Since they define transitions between items as being completely deterministic and based on the largest representational similarity between them, trajectories always enter a loop. Given such simple transition rules, the relation 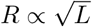 can be derived, where *L* is the number of items to recall (a similar derivation can be found in Appendix).

In contrast, the dynamics in our model are not deterministic (we will discuss this point in Section 5), and we hardly ever observe a loop in the network trajectories. However we can still compute a similar measure to *R*, labeled as *M*_it_, which is the number of retrieved patterns until the network repeats one transition. Compared to the measure *R* computed in [6], our measure *M*_it_ has a steeper slope, though again lower than 1 (Fig. 2a). Thus, *M*_it_ still grows sublinearly with respect to *L* but it does not match the *R* measure, whose slope is 0.5. Alternatively, we have computed the number of retrieved items until the network simply revisits one of the already-visited items, labeled as *M*_i1_. In contrast to *M*_it_, this latter quantity shows a slope *lower* than 0.5, which again does not match the *R* trend in [6] (Fig. 2a). For this reason, we define a new quantity called *M*_i_(*L*), which is the number of recalled items until one item is repeated *twice*. This somewhat contrived quantity has a behaviour indeed similar to that theoretically expected from the quantity *R*, that is, a slope of 0.5 (Fig.2b).

**Figure 2:**
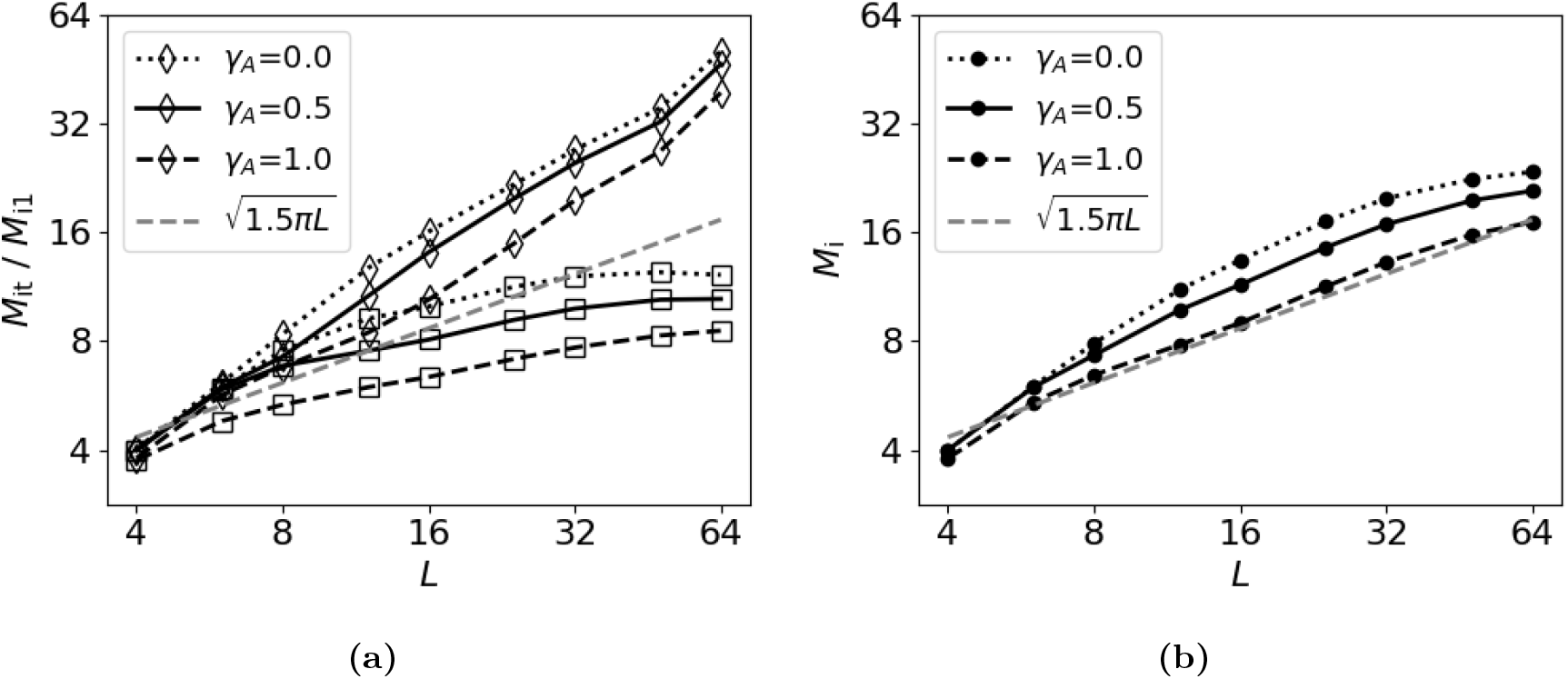
A repetition-limited measure of performance in free recall approaches a 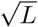 dependence. The dashed gray line is the theoretical prediction of *R* in [6]. Both axes are in a log scale. **(a)** *M*_it_ is the number of recalled STM items until one transition is repeated (marked by diamonds). *M*_i1_ is the number of recalled STM items until one of the visited STM items is revisited (marked by squares). Dotted curves are from the slow inhibition regime (*γ _A_* = 0.0), solid curves are from the intermediate regime (*γ_A_* = 0.5) and dashed curves are from the fast inhibition regime (*γ_A_* = 1.0). **(b)** *M*_i_, the number of recalled STM items until one of them is repeated *twice*. In contrast to the two measures plotted in (a), this quantity approaches a square root dependence with *L*.

In computing these three measures, we have ignored errors (extra-list items) in order to compare with [6, 11]. Note that errors are not discussed in their conceptual model and experiment, in which retrieval of extra-list words is simply dismissed as irrelevant. The beauty of their treatment, in fact, stems from the simple question they pose, without getting into how the recall process happens dynamically in the brain and how LTMs affect performance of free recall. These questions are our own interest in this work.

Moreover we find that the three inhibition regimes considered here (slow, fast and intermediate regime, see Methods) behave similarly in terms of short-term memory function. Based on this observation, hereafter we only concentrate on the intermediate regime (*γ_A_* = 0.5).

### 3.2 If limited by duration, the network *can* again recall up to 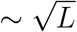 STM items

In the free recall experiment conducted in [6], they computed *R* as the number of correctly recalled words (or sentences), ignoring errors and repetitions. The time allocated to recall started from 1 minute and 30 seconds for *L* = 4, and was increased by the same amount when the length of the list was doubled. As it is problematic to establish a correspondence between human recall time and simulation time in the Potts model, we define another quantity: we compute the number of correctly retrieved items, ignoring errors and repetitions, *M*_u_, within a *given number of consecutive latches*, denoted by *g*(*L*). Given the stochasticity of the network dynamics in visiting pattern space, the specific choice of *g* (*L*) has implications on *M* _u_. Therefore, we set *g*(*L*) = 4 log_2_(*L*) − 2 in order to establish a reasonable comparison with the results in [6]. We find that this measure has a slope of approximately 0.5 (Fig. 3). However, if *g*(*L*) = *L*, i.e., a linear function of *L*, *M*_u_ has a higher slope. Finally, if we set *g*(*L*) to *g*(*L*_max_) = 22, with *L*_max_ ≡ 64, i.e. constant and equal to the maximum number of latches in the logarithmic option, then *M*_u_ becomes slightly larger for intermediate values of *L*, suggestive of a drop after hitting a maximum. This again indicates that the Potts model can capture the empirical trend of 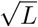, provided one adopts a suitable rule for limiting the length of latching sequences.

**Figure 3:**
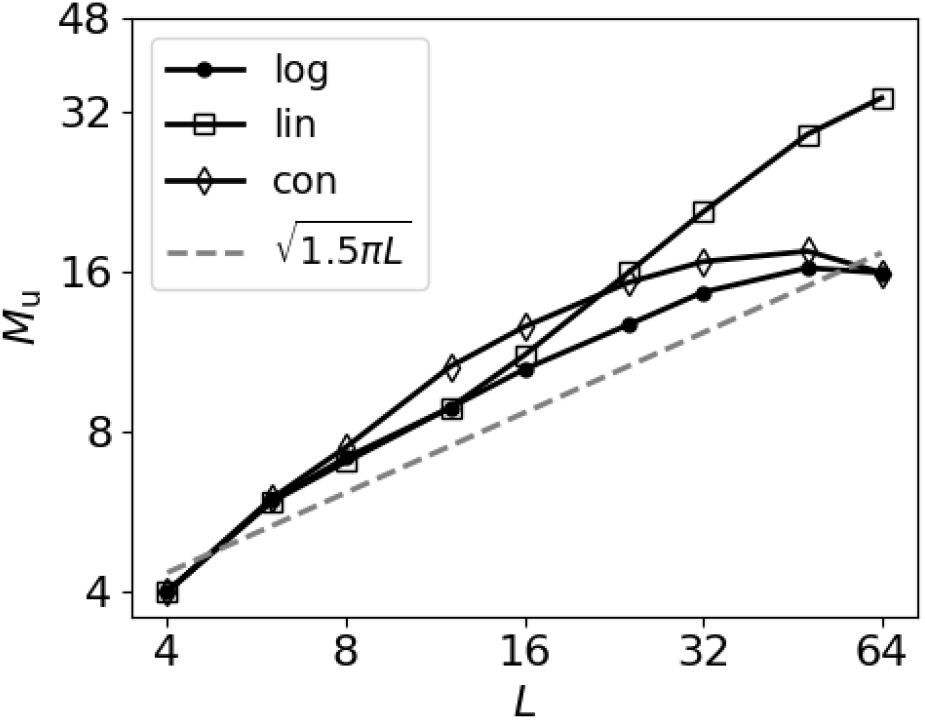
A duration-limited measure of free recall can also approach **a** 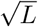 dependence. *M*_u_, the number of recalled STM items within a given number of latches, *g*(*L*), is plotted as a function of *L* in log-log scale. We consider three different functions for *g*(*L*): logarithmic, linear and constant function of *L* denoted by dots, squares and diamonds, respectively. The dashed gray line is the theoretical prediction of *R* in [6].

### 3.3 Free recall of locations in a 2D grid also shows an approximate 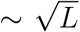 dependence

That the various *M* measures obey quasi-square-root functions of *L* may be partially understood by considering a random walk in pattern space, with equally probable visits to each of the patterns (Fig. S6) [12]. Inspired by this observation, we have designed simple experiments in which subjects are asked to remember a random trajectory on a 2-dimensional grid (Fig. 10). We then asked participants to freely recall the positions of the presented dots by clicking on their positions on the grid.

Clearly, the parameters of the experimental protocol can be expected to affect recall, including the amount of time allocated for recall. However, in our experiment, participants only need to click on the correct locations (as opposed to typing in the words they recall [6]), and setting a fixed recall time may seem *ad hoc*. As an alternative, and to further explore the validity of latching dynamics as a model for this experiment, we give participants a limited number of clicks per trial, set as 2(*L* − *h*(*t*|*L*)), where *h*(*t*|*L*) is the number of correctly recalled dots up to that point in time. Then we computed *M*_R_, defined as the number of correctly recalled dots for a given *L* ignoring errors and repetitions, and compute the same measure from simulations with the Potts model (see Methods for a detailed description of the experiment as well as how to compute *M*_R_).

We find a reasonable agreement between the performance of the Potts model and human subjects in our experiment, where both cases show a slope of approximately 0.5 (Fig. 4). This suggests that latching dynamics capture some aspects of the underlying neural mechanisms of free memory recall, perhaps related to the random walk nature of the trajectory, although the exact details depend on the paradigm.

**Figure 4:**
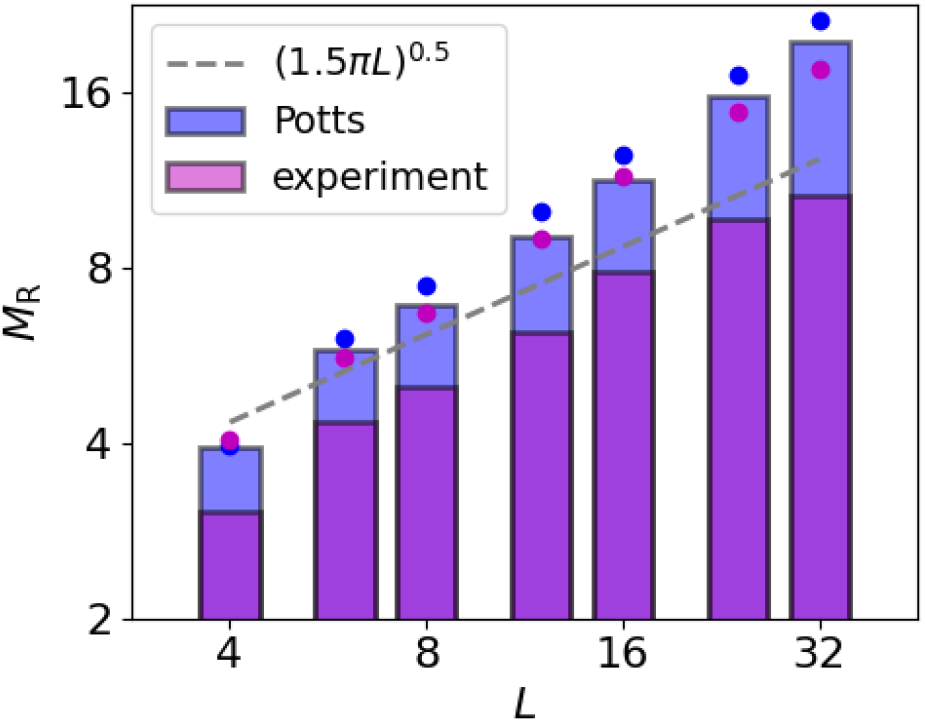
Free recall of locations in a 2D grid also shows an approximate 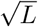 dependence. *M*_R_, the average number of correctly recalled locations in our experiment, is shown by the height of pink bars in a log-log scale. The distance from the bar to the dot of the same colour corresponds to the standard deviation of the mean. Results of 40 participants are pooled together. The same quantity *M*_R_ is computed, from simulating Model 2, as the number of correctly retrieved STM items within a given number of consecutive latches set as 2(*L* − *h*(*t*|*L*)), where *h*(*t*|*L*) is the number of correctly recalled STM items up to that point in time (blue bars). The dashed gray line is the theoretical prediction of *R* in [6]. Both results, from our experiment and the Potts model, show an approximate 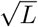 trend.

### 3.4 If limited by errors, the network cannot recall beyond its STM capacity

The measure *M*_corr_ was introduced and discussed in section 2 for comparing three different models. Here we compute the same quantity with a slight modification; in order to compare with our experimental data, we consider sequences of variable length that depends both on list length *L* and time. We consider lengths of *g*(*L*) = 2(*L* − *h*(*t*|*L*)), where *h*(*t*|*L*) is the number of correctly retrieved STM items up to that point in time; within this sequence we count the number of correctly recalled STM items until there is either an error or a repetition. We compute this quantity 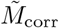 for several values of Δ*θ* in the Potts model. We find that the behaviour of 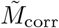 with respect to *L* is qualitatively similar to that of the experimental curve for a broad range of Δ*θ* values (see Fig. 5a). For all values of Δ*θ*, 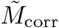 saturates reaching a maximum that is similar to that of the experimental data, of around 8 items correctly recalled. Exceptions are at the two extremes: too small and too large values lead to lower capacity of the Potts model, below 7 items.

**Figure 5:**
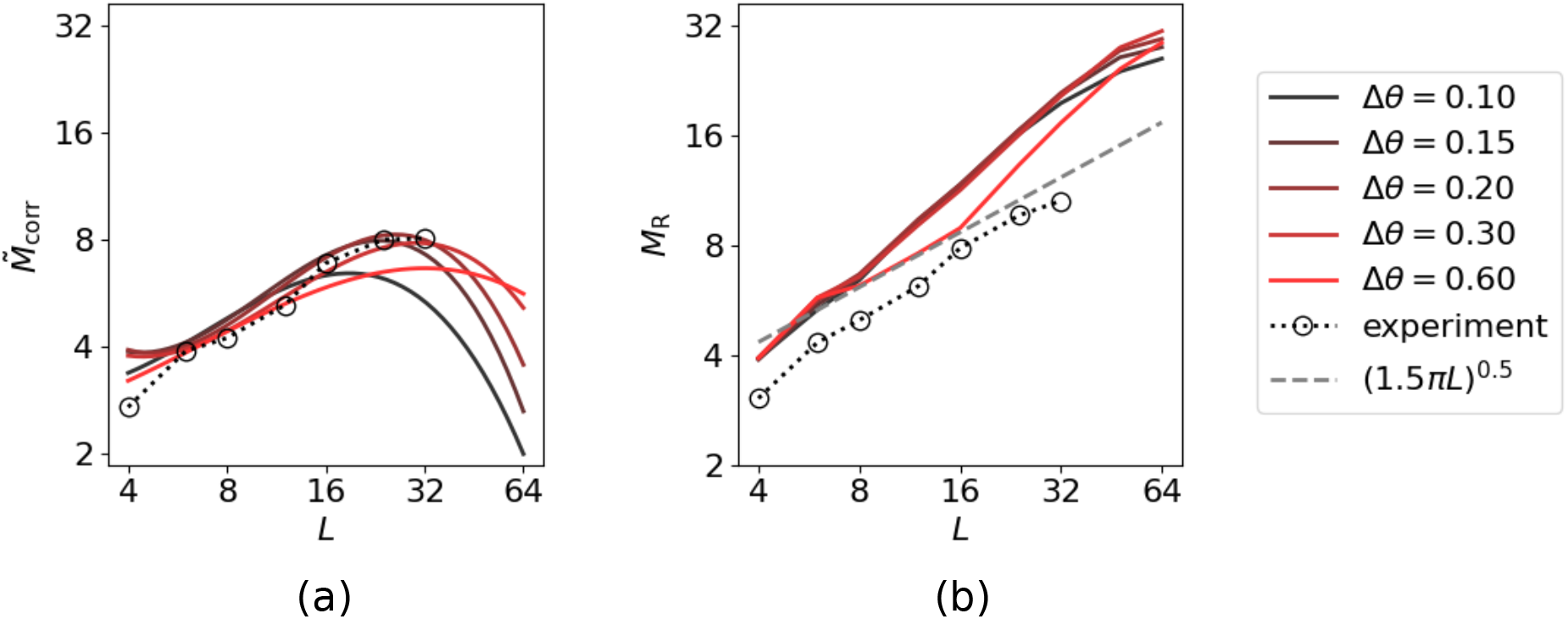
An error-limited measure of recall has a maximum value. Two measures, 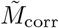 and *M*_R_, are shown for several values of Δ*θ*, coded by colours. Black dotted curves are the experimental results of free recall of locations in a 2-dimensional grid. **(a)**: 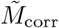 has a maximum value. It is the number of recalled STM items until the network either revisits one of the already-recalled STM items or visits one of the LTM items, but within a given number of latches – 2(*L* − *h*(*t*|*L*)), where *h*(*t*|*L*) is the number of correctly recalled STM items up to that point in time. **(b)**: *M*_R_ shows a scaling behaviour. *M*_R_ is the number of recalled STM items, ignoring repetitions and errors, within a given number of consecutive latches set as 2(*L* − *h*(*t*|*L*)), where *h*(*t*|*L*) is the number of correctly recalled STM items up to that point in time.

The saturation behaviour, and hence the notion of memory capacity, again contrasts with the scaling behaviour approximated by the various measures such as *M*_i_, *M*_u_ and *M*_R_. This contrast holds irrespective of the values of network parameters used in simulations. Indeed the scaling behaviour of *M*_R_ is almost independent on the value of Δ*θ* except when it is too large, Δ*θ* = 0.6 (Fig. 5b). Furthermore, we find that the two contrasting behaviours – scaling and saturation – are fairly robust to change of network parameters such as Δ*θ*, *S* and *a* (Figs. S7 and S8).

“Performance” therefore depends very differently on *L*, if recall is taken to be terminated by errors, i.e. by the recall of an item that is not in STM. Thus, while if ignoring errors the notion of STM capacity appears irrelevant (evident from the scaling behaviour of the various quantities discussed above), it becomes relevant if errors are considered to be critical in the task.

In summary, we have shown in this section that whether we get scaling or saturation in the performance depends on the specific metric we use to measure the performance, both in the Potts model and in our experiment. In free recall experiments, performance has often been quantified through the *M*_R_ index, thereby ignoring errors. The scaling behaviour of this index has recently been corroborated for lists up to 512 words [6]. In contrast, taking our experiment as an example, we have shown that if errors are considered critical, in our case through the *M*_corr_ measure, then the performance of human subjects actually expresses a saturation at about 8 items. In our model, that expresses a similar behaviour, this saturation is brought about by the interference from long-term memories.

## 4 Serial recall

We have found that repetition-limited and duration-limited measures of performance in free recall in the Potts model endowed with short term memory function can express quasi-square-root behaviour in the number of items in the list. One question that naturally arises is whether the same model can express behaviour similar to *serial recall*, a paradigm very similar to free recall, but with a crucial difference. Here, participants are instructed to recall items in *the same order* as they have been presented, making the task more difficult.

We have performed serial recall experiments with three different types of materials. We asked the participants to observe and repeat sequences of stimuli presented to them on the screen - either digits or spatial locations on a 2-dimensional grid (Fig. 10), and varied the time of presentation of the stimuli in the observed sequence. There were two conditions for the spatial locations, referred to as Locations and Trajectories: in the Locations condition, considered to involve only “discrete” items, the six chosen locations around the centre of the grid were highlighted in any order, while in the Trajectories condition, every next location was one of the six consecutive locations around the previous one, thus suggesting a “continuous” trajectory. Contrary to the previous experiment reported in Section 3.3, in this task participants had to recall the material in the correct order, otherwise the trial was dismissed as incorrect. Participants started with short sequences of length 3; if they recalled them correctly in at least 3 out of 5 trials, the sequence length increased, until a memory capacity limit for this stimulus type and presentation time was reached. In this way we measure the memory capacity for serial recall by computing the *Area Under the Curve* (Fig. 6).

**Figure 6:**
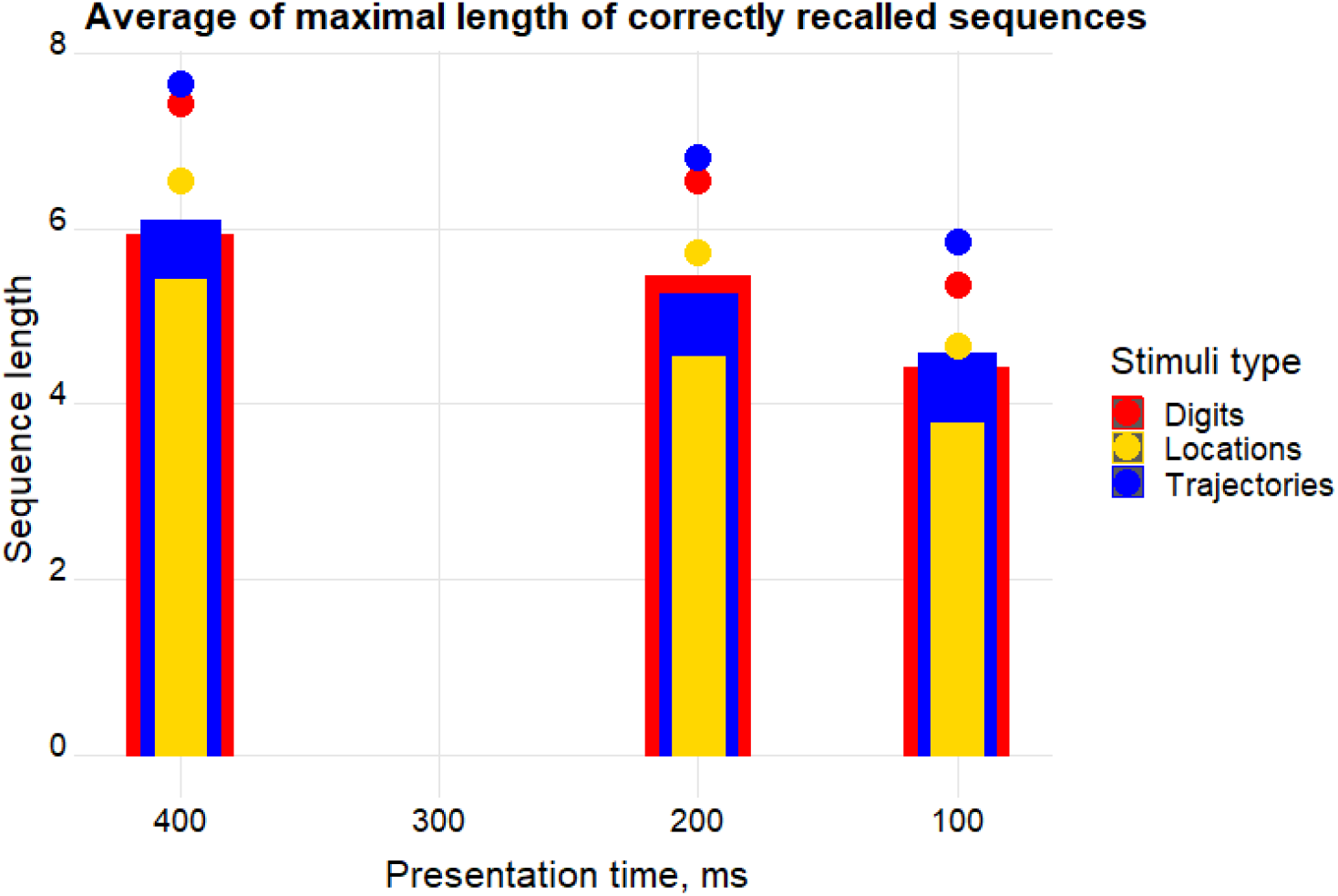
Short-term memory capacity for serial recall does not markedly depend on stimuli type. Memory capacity for serially presented stimuli for different presentation times: bars correspond to the average across participants of the longest correctly recalled sequence, while the distance from the bar to the dot of the same colour corresponds to the standard deviation of the mean. We performed the experiment for three different stimulus types, shown in different colours.

Our experiment yields two main results (Fig. 6). The first is that the type of stimulus does not affect the recall probability, except for a slight disadvantage in the *discrete* Locations condition, suggesting a universal mechanism for recall independent of the material, which manifests itself at the systems level. The second, which is more pronounced, is the effect of presentation time per stimulus, that, when shortened, makes it more difficult to correctly remember and repeat the longer sequences, suggesting a disadvantage at the *encoding* stage. We ask whether latching dynamics in the Potts model can express this finding. Given that our results, as well as those from other studies [7], show very little dependence on stimulus material, hereafter we only consider the result with digits in order to establish a comparison with our model.

We used Model 2 (lower adaptive threshold for items held in STM) to constrain the dynamics into a subset of *L* = 6 patterns intended as the 6 digits of our experiment. In addition to that, we introduced heteroassociative weights, similar to Model 3, to provide the sequential order of presented digits (see Eq. (13)).

We find a good agreement between our experimental data and the model (Fig. 7). In addition, we find that human subjects perform better if the to-be-memorised digit series include ABA or AA (Figs. 7a, 7c), in line with the notion that the repetition of an item aids memory [13, 14, 15, 16]. Such sequences are not produced by our model, due to firing rate adaptation and inhibition preventing the network from falling back onto the same network state for time scales of the order *τ*_2_. Due to this, we refer to stimulus sets *excluding* such sequences as *Potts-compatible*.

**Figure 7:**
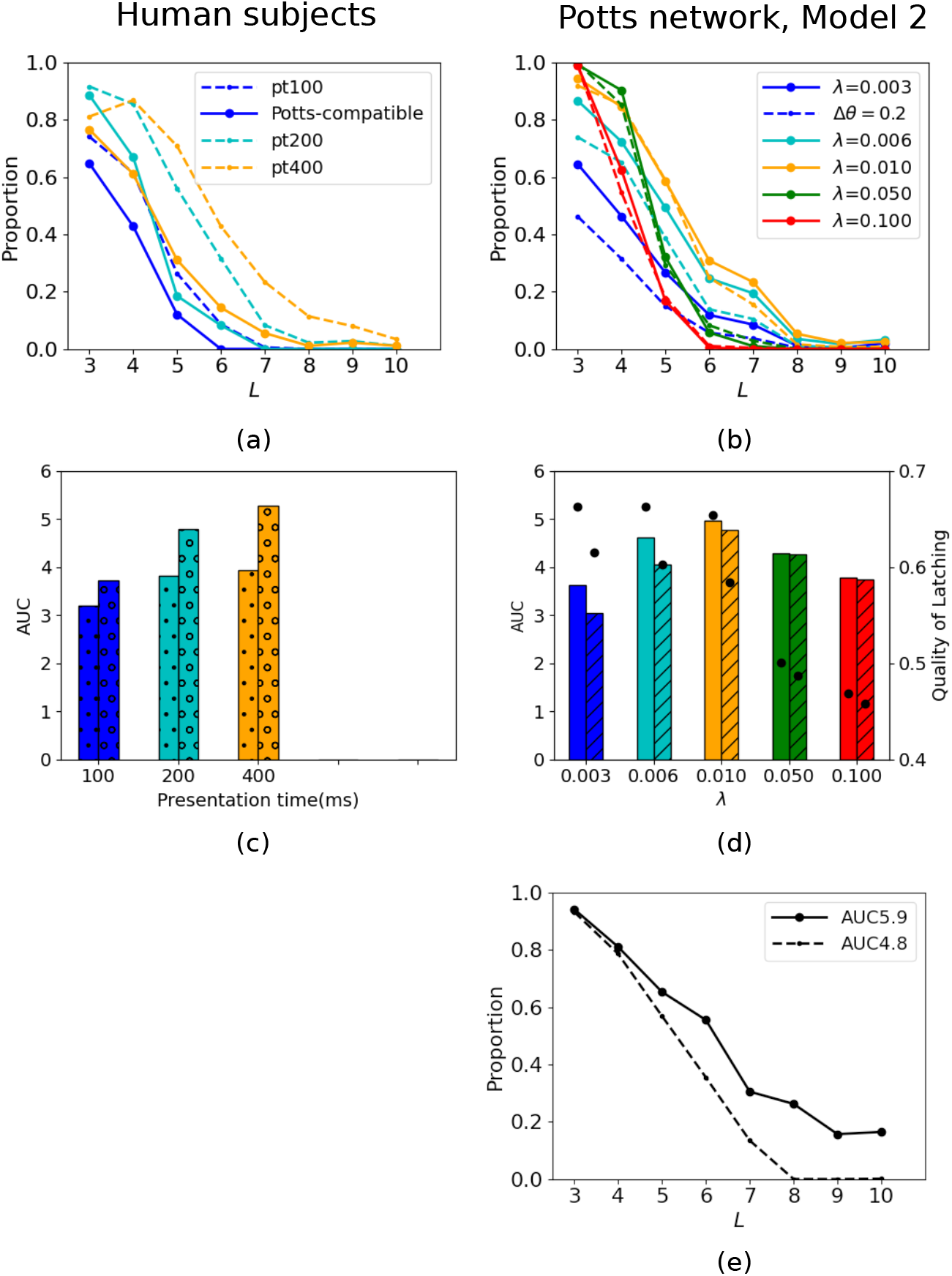
Serial recall of digits by human subjects and the Potts model. **(a)** Proportion of correct trials in the serial recall task with digits. Data for all subjects (*n* = 36) are pooled together. Colour codes for presentation time in units of milliseconds. Solid and dashed curves for each colour show the result of “Potts-compatible” (see Methods) trials and of all trials, respectively. **(b)** Proportion of correct subsequences in a latching sequence of the Potts model. Colour codes for values of *λ*. Solid (dashed) curves are for Δ*θ* = 0.1(0.2). **(c)** *Area Under the Curve* (AUC) computed from the curves of (a). Colour-coding is the same as in panel (a) and the bars filled with dots (open circles) correspond to solid (dashed) curves in (a). **(d)** AUC for latching sequences of the Potts model. Same colour-coding as in (b) is used. Bars without hatches are for solid curves in (b) and those filled with oblique lines are for dashed curves in (b). Black dots indicate the quality of latching on the right y-scale. **(e)** Proportion of correct subsequences in a latching sequence of the Potts model. The solid curve is for congruent instructions only and the dashed curve is for a shuffled version of intrinsic sequences.

The heteroassociative component of the learning rule (Eq. (13)) provides “instructions” to the network regarding the sequential order of recall, allowing it to perform serial recall (this is to be contrasted with the model with a purely autoassociative learning rule, performing free recall). The strength of such instructions is expressed through the parameter *λ*. We find that this parameter plays a role similar to that of presentation time in our experiments; increasing it enhances performance, just as increasing the presentation time increases the performance of human subjects (Fig. 7). However, values of *λ* that are too large again make performance worse and deteriorate the quality of latching (Fig. 7d). The dynamics is a stereotyped sequences of patterns, see Fig. S9, without any nonlinear convergence towards attractors, and the sequence itself is progressively harder to decode. Therefore, the most functional scenario is when the heteroassociative instruction acts as a bias or a perturbation to the spontaneous latching dynamics rather than enforcing strictly guided latching in the Potts model. This is in sharp contrast with the mechanism for sequential retrieval envisaged in the model considered in [17], where the heteroassociative connections are the main and only factor driving the sequential dynamics; in that case, without it, there are no dynamics but rather, at most, the retrieval of only the first item. The effect of lower adaptive threshold (expressed by Δ*θ*) on latching sequences is to constrain the dynamics to a subset of presented items among *p* patterns, but values of Δ*θ* that are too high degrade the performance as well as the quality of latching (Fig. 7b, 7d).

As mentioned above, the Potts model produces latching sequences even without any heteroassociative instructions. This means that the free transition dynamics of the model may or may not coincide with the “instructions” provided by the heteroassociative weights. Then one question naturally arises. How does the congruity between spontaneous, endogenous sequences and instructed ones affect the performance of the model? To see this effect, we obtain some intrinsic sequences by running simulations with *λ* = 0; from these sequences, we generate a set of instructions. These instructions are *congruous*, as they reproduce latching sequences emerging without any heteroassociative instructions. Then we compare the performance for these congruous instructions with those of incongruous instructions, which we obtain by shuffling the congruous ones. We find that the capacity of the model (denoted as AUC in the legend in Fig. 7e) increases by as much as 1 item for the congruous case relative to the incongruous case.

These results together with those from the previous two sections indicate that intrinsic latching dynamics, similar to a random walk, can serve short-term memory (e.g., free recall). Furthermore latching dynamics can also serve serial recall, if supplemented by biases that modify the random walk trajectory; the modification (or perturbation) should be a quantitative one, which biases the random walk character of the trajectories, rather than an all-or-none, or qualitative one, that inhibits it. This is consistent with our recent experimental result, where “guided” serial recall leads to poorer performance than a non-guided control (O. Soldatkina, in preparation).

## 5 The trajectories in free recall

In previous sections we saw a reasonable agreement between some experimental measures and those extracted from simulating the Potts model. This agreement essentially results from two factors: first, the Potts model can produce a sequence of discrete activity patterns even though its governing equations are continuous at the microscopic level; and second, the dynamics of the Potts model visit the patterns in a random-walk like process. We now examine the sequences more closely to see what factors influence latching sequences and how the network wanders around the landscape of memorized patterns.

We first ask ourselves: once the network is cued with a given pattern, what elicits the retrieval of the next one? In previous studies [18, 9], it was shown that transitions occur most frequently between highly correlated patterns, when the Potts model serves a long-term memory function. We confirmed that this is also the case when the Potts model serves a short-term memory function, as in the current study (Fig. S10). Indeed, the larger the average correlation of one pattern with all other patterns in STM, the more often it is visited by the network (Fig. S11). This result is consistent with a recent experimental study on how memorability of words affects their retrieval in a paired-associates verbal memory task [19].

Next we probe the flow of information in the latching sequences of the STM model embedded in the Potts neural network by computing the normalized mutual information between two patterns as a function of their relative separation in a latching sequence, *z* (Fig. 8a). We find that the mutual information is decreasing rapidly with respect to *z*, with a quasi-periodic modulation, reminiscent of the temporal profile of intensity of a damped oscillator. The periodic modulation is much more evident for *L* = 16 than for *L* = 64; within the range of *z* we have considered, we see a peak at *z* ≈ 4.5 for *γ_A_* = 0.0 and at *z* ≈ 3.5 for *γ_A_* = 0.5, but we also see the *second-order* peak at *z* = 6 in addition to the *first- order* peak at *z* = 3 for *γ_A_* = 1.0 (Fig. 8a). The second-order peaks for *γ_A_* = 0.0 and *γ_A_* = 0.5 would be located at *z* ≈ 9 and *z* ≈ 7, respectively. The quasi-period of the “damped oscillation”, *ζ*, is twice the *z*–value of the first peak, therefore, decreasing with increasing *γ_A_*, starting from *ζ* ≈ 9 at *γ_A_* = 0.0 until *ζ* ≈ 6 at *γ_A_* = 1.0. For *L* = 64, it’s as if the damping ratio is too high to observe any periodicity.

**Figure 8:**
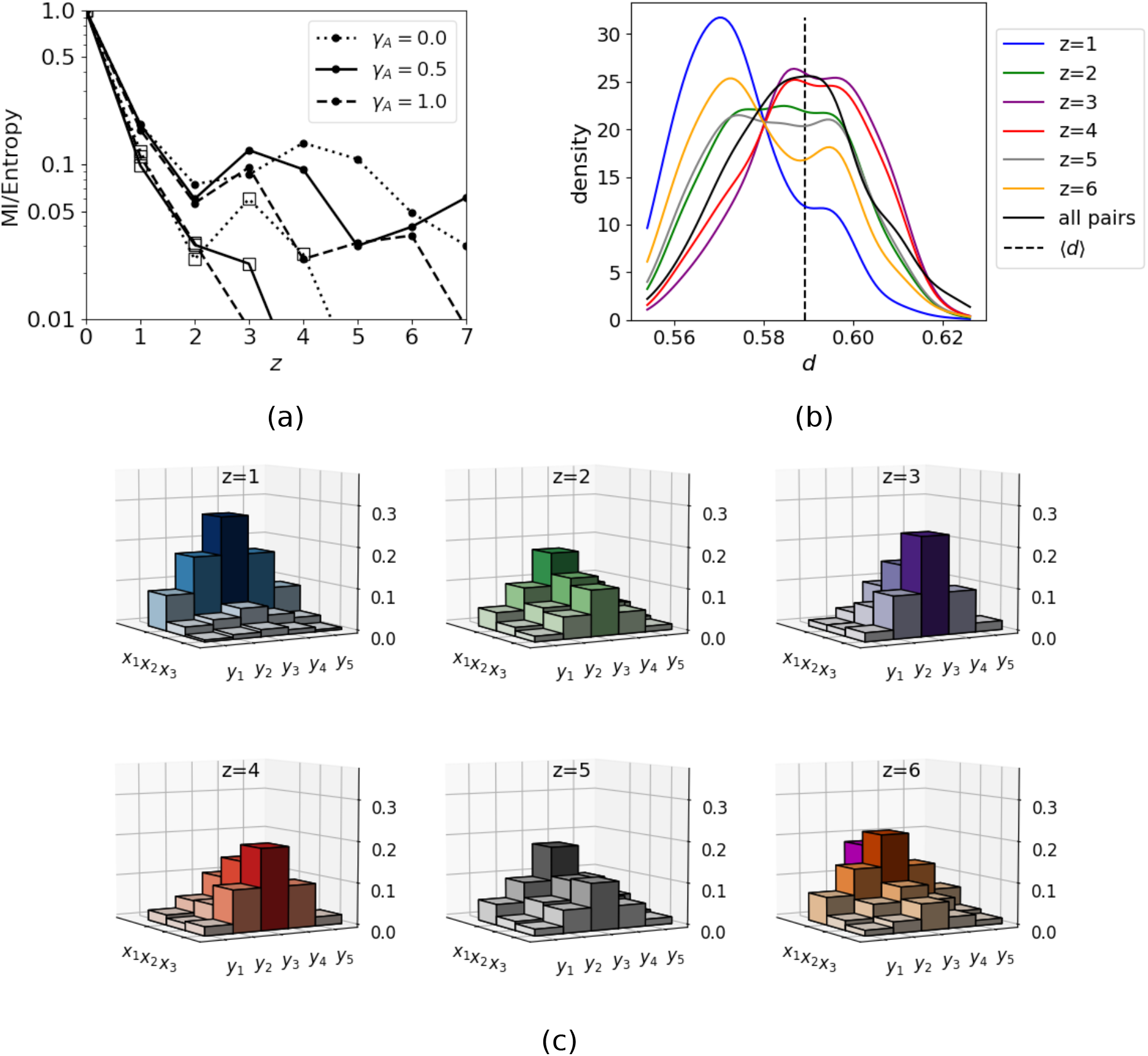
Damped waves in pattern space. **(a)** Mutual information as a function of the relative separation of two items, *z*. The ordinate is the mutual information *I*(*μ_n_, μ_n_*_+*z*_) divided by the entropy *H*(*μ_n_*) ≡ *I*(*μ_n_, μ_n_*). Note the logarithmic scale of the *y*–axis. Parameters are Δ*θ* = 0.3, *L* =16 (64) for the curves marked with dots (open squares), *w* = (0.4, 0.8, 1.0) for *γ_A_* = (0.0, 0.5, 1.0). **(b)** Distribution of distance, *d*, between two patterns that have the relative separation *z* in a latching sequence for *L* = 16, *γ_A_* = 0.5 and *w* = 0.8. The black, vertical line indicates the mean value of *d* across all *p* patterns. The solid black curve is the PDF of *d* among all possible pairs between L patterns in STM. **(c)** Histograms for the visiting frequency of patterns in STM, given one pattern is recalled. The remaining *L* − 1 = 15 patterns are arranged along the *x*–axis by their visiting frequency at the next position of the currently retrieved pattern in a sequence (*z* = 1), giving three groups *x*_1_, *x*_2_ and *x*_3_ of 5 patterns each. Each group is further arranged symmetrically along the *y*–axis, with the most frequent pattern on the midline (*y*_3_). Visiting frequency is double-encoded by the height and colour of bars. The lonely, magenta bar behind the group *x*_1_ shows the visiting frequency of the currently recalled pattern once it returns at the position *z*.

This behaviour is related to how the Potts network “freely” forages the landscape of the embedded attractors. We visualize this nontrivial behaviour for *γ_A_* = 0.5, where we not only see a kind of damped *wave* that “propagates” along the *y*−axis with the variable *z* as an effective “time”, but also see the “reflection” of the wave around *z* ≈ 3.5 (Fig. 8c).

What causes these characteristics of the latching trajectories of the Potts model? To answer this question, we define a quantity, called *d*, which is an index of “semantic” distance between two patterns in their representational space. We defined a distance between two patterns *μ* and *ν* as follows.

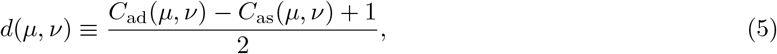

where *C*_as_ and *C*_ad_ measure the correlation between two patterns (see Eqs. (**??**)).

We consider the distribution of *d*(*μ_n_, μ_n_*_+*z*_) for 6 values of *z* (Fig. 8b). At *z* = 1, latching occurs mostly between highly correlated patterns as expected, where the higher correlation is expressed by lower *d*. At the second step in a latching sequence (*z* = 2), patterns that have higher *d* values than the average value 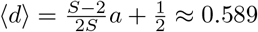 show a comparable proportion of the probability density curve relative to patterns with lower values of *d*. Then the proportion of higher *d* values is much larger than the proportion of lower *d* values for *z* = 3 and *z* = 4. This means that the network prefers to visit those patterns that are less correlated with the initially retrieved one at the third and fourth step. So we can say that the network reaches the most “distant” pattern from its “initial” pattern around *z* = 3.5, which is the “reflection” point of the wave (Fig. 8c). As *z* increases further to reach 6, the density curve is getting closer to the curve for *z* = 1, thus approaching the periodicity mentioned above. This periodicity is confirmed by Figs. S12, S13.

These results indicate that latching trajectories by Potts networks have a quasi-random walk character, though biased by correlations between patterns in their representational space. This is consistent with earlier applications of latching dynamics to semantic priming [5].

## 6 Discussion

The Potts model offers a plausible cortical framework to discuss aspects of memory dynamics, without losing too much of the clarity afforded by simpler non-neural models. Indeed, a major difficulty with network models of memory storage in the human cortex, which have attempted to reflect its dual local and long-range connectivity [20, 21] by articulating interactions at both the local and global levels, is that their mathematical or even computational tractability usually has required *ad hoc* assumptions about memory organization. For example, the partition of memory items in a number of classes, in each of which memories are expressed by the activity of the same cortical modules [22] – which makes it awkward to use such a network model to analyse the free or serial recall of arbitrary items. On the other hand, more abstract models have provided brilliant insight [11] which is hard, however, to relate to neural variables and neural constraints. By subsuming the local level into the dynamics of individual Potts variable, the statistical analysis can focus on the cortical level, what is effectively a reasonable compromise.

The (global) cortical level is in particular the one to consider in assessing short-term memory phenomena, in which interference from widely distributed long-term memories plays a central role. Experiments with lists of unrelated words are a prime example [5]. The free energy landscape of the Potts model provides a setting for quasi-discrete sequences of states, with properties that turn out to be similar to those of random walks. This happens, however, only within a specific parameter range, and only to a partial extent, so that often one has in practice several intertwined sequences, with simultaneous activation of multiple patterns, as well as pathological transitions, all characteristics with potential to account for psychological phenomena, and which are lost in a more abstract purely symbolic model. We have thus discussed three generic neural mechanisms that may contribute to restrict the random walk, approximately, from *p* to *L* items. Although not exclusive, we have argued that the second such mechanism is the one most relevant to account for the recall of list of unrelated items.

To model the recall of ordered lists, an additional *heteroassociative* mechanism can be activated, which biases the random walk, but again approximately, resulting in frequent errors and limited span. We have observed that, at least in the Potts network, if the heteroassociations, which amounts to specific instructions, dominate the dynamics, the random character is lost, but with it the entire latching dynamics – which cannot be harnessed to just passively follow instructions.

In summary, a Potts network can generate quasi-digital sequences from analog processes, with the possibility of errors in

1. the digitalisation into a string of discrete states, one at a time
2. the restriction to *L* out of *p* item in LTM
3. the order, both in the specific sense of serial order, and in the generic one of avoiding repetitions.

These possibilities for error reflect weaknesses of latching dynamics as a mechanism for short-term memory expressed by a Potts network, and at the same time underscore the value of the mechanistic model, inasmuch as similar “flaws” crop up in the phenomenology. The analysis of such flaws can lead to refinements of the model.

Thus, point 2, the difficulty of restricting latching dynamics to a subset of all the long-term memory representations, is made even more severe in paradigms that involve multiple subsets. If one accepts the evidence of a common substrate for working memory and long-term memory representations [23], one cannot resort to different “drawers”, i.e., different scratchpads or the like, where to temporarily hold the items from distinct subsets, and this makes enforcing the restriction more difficult. Likewise, one cannot regulate the correlation between the long-term representations, as one could do if new *ad hoc* representations were temporarily set up. These constraints can result in intrusions, a simple form of false memory, e.g. by items that are strongly semantically associated to items in a short-term memory list [24], or by items in prior lists [25]. It would be tempting to pursue a fully quantitative study of these phenomena [26] to try and extract constraints, for example, on the time course of the “kick” in the Potts model.

In relation to point 3, latching dynamics are intrinsically stochastic in nature, even in the absence of microscopic noise, because of the heterogeneity of the underlying microscopic states. With randomly correlated representations, trajectories among items are effectively random, with only a tendency to avoid close repetitions, as a result of the adaptation-based mechanism. Interestingly, a tendency to perceive random processes as less prone to repetition than they really are is a hallmark of human cognition [27]. Beyond the vanilla version of the model, however, it is rather trivial to incorporate e.g. adjustments of the time course of the kick, to produce primacy and recency, or adjustments of the correlations between pairs of representation to produce preferred transitions. What is more interesting and still lacking, to our knowledge, is again a quantitative study of the degree of randomness of the recall process, in the context of remembering lists for example – a study made inherently difficult by the need to use novel items in a within subjects design. The same need effectively prevents the analysis of the recalled string at the single neuron level: even when recording the activity of neurons in awake patients, only generic forms of selectivity can be reliably studied, e.g., that expressed by putative “time” cells [28]. Lacking experimental evidence on the dynamics of individual neurons, the Potts model can help interpret evidence at the integrated cortical level.

It is its fallibility in the production of a simple string of items, however, where the Potts network offers crucial insight beyond that provided by simpler and more abstract models, in which the digitalisation of a string is *a priori* given. Latching dynamics can involve partially parallel strings, items incompletely recalled simultaneously with others, periods of utter confusion, stomping attempts. Statistically, they are all observed with prevalence determined by the various parameters. These flaws in the analog-to-digital transduction of the Potts model may be useful in the interpretation of electrophysiological data. One basic question in this domain is: can two items be simultaneously active in working memory? On this question, experimental evidence has been difficult to obtain, because a process that appears to involve two items active together, might in fact rapidly alternate between them. Recently, however, the genuinely concurrent activation of two items has been reported with a model-based analysis of EEG data [29]. In that study, holding on to the two items meant better performance in the task, so it reflects a capability, not a flaw of the short-term mechanism. If extended to sequences of endogenously generated states, as the Potts model indicates would occur, at least in certain regimes, it would mean that not only the focus of attention when performing a similar task need not be unique, but also that parallel streams of thoughts can be entertained along partially interacting trajectories. This could be applied to interpret electrophysiological measures of mind wandering dynamics [30], with significant implications for our intuition about the unity of consciousness [31].

## 7 Model and Methods

### 7.1 Potts neural networks

A Potts neural network is an autoassociative memory network [32], comprised of *N* Potts units, which can be activated along one of *S* states or remain in a quiet state. The *N* units interact with each other via tensor connections, whose values are pre-determined by a Hebbian-learning rule,

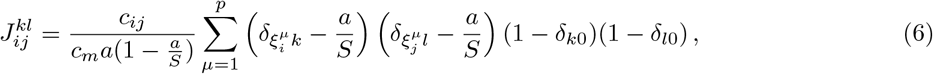

where *c_ij_* is either 1 if there is a connection between unit *i* and *j* or 0 otherwise. The average number of pre-synaptic units for a given unit is 〈*c_ij_*〉 = *c_m_*. We use randomly-correlated memory patterns 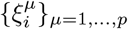 in this work but one can consider a set of correlated memory patterns, as produced by the algorithm presented in [33]. Sparsity of patterns (fraction of active units) is set by the parameter *a*.

As the Potts model is considered as a model of cortical dynamics, the threshold component 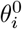 specific to unit *i*, but acting on *all* its active states, varies in time with its own time course, associated in previous papers to a single time constant, *τ*_3_. This threshold is intended to effectively describe local inhibitory effects, which in the cortex are relayed by at least 3 main classes of inhibitory interneurons [34] acting on GABA_A_ and GABA_B_ receptors, with widely different time courses, from very short to very long.

The time evolution of the network is governed by the equations

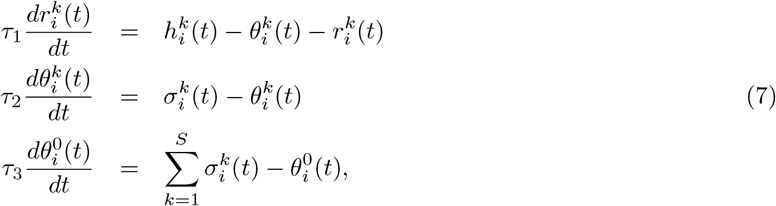

where the variable 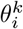 is a specific threshold for unit *i*, specific also for state *k*, varying with time constant *τ*_2_, and intended to model adaptation, i.e. synaptic or neural fatigue selectively affecting the neurons active in state *k*, and not all neurons subsumed by Potts unit *i*. The “current” that the unit *i* in state *k* receives is

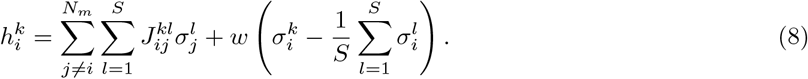

Here *w* is another parameter, the “local feedback term”, first introduced in [9], to model the stability of local attractors in the full model, as justified with a semi-analytical derivation later [8]. It helps the network converge towards an attractor, by giving more weight to the most active states, and thus it effectively deepens the attractors.

The activation along each state is given by

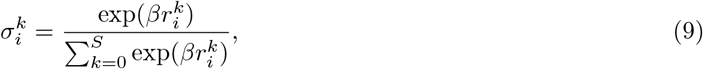

where 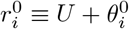.

#### Combining slow and fast inhibition

Latching dynamics, in the model equipped with either slow or fast inhibition, are studied in [9, 18]. Here, we consider a more realistic case in which *both* slow and fast inhibition are taken into account. Formally, it is based on replacing the inhibitory or non-specific threshold 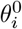 with the sum 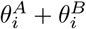 (to denote fast, GABA_*A*_ and slow, GABA_*B*_ inhibition, respectively) and writing separate equations

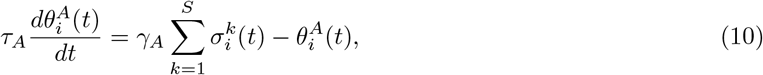

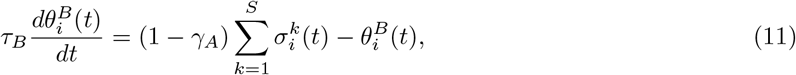

where, instead of *τ*_3_ either short or long, one sets *τ_A_ < τ*_1_ *<< τ*_2_ *<< τ_B_* and *γ_A_* as the parameter setting the balance of the two. Accordingly, 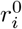 should become 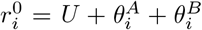 in Eq. (9). Note that a more realistic approach would be to consider inhibition at all time scales, in line with experimental findings [34]. By introducing very fast and very slow inhibition, we only make a small step towards plausibility, while maintaining relative mathematical simplicity, and possibly being able to apply separation of time scale methods to better understand the phenomenology.

#### Phase diagram of latching dynamics

We studied different latching behaviours of Potts network by extensive computer simulations. In the simulation the network is first initialized by setting all variables at their equilibrium values. Then we cue the network with one of the memorized patterns and let the dynamics proceed. Simulations are terminated if the network is trapped into an globally stable null attractor (where all units are inactive) or if the total number of updates reaches 10^5^.

The Potts network has four different phases in the parameter space considered here (Fig. 9). The first one is the trivial *no-latching* phase, where the network dynamics stop without a retrieval of patterns other than the cued one. The second one is what we are interested in as a model for short-term memory, i.e., the *finite-latching* phase, where the quality of latching is good. The third phase is *infinite-latching* one, where the dynamics go on endlessly once being cued. In the fourth phase the retrieved pattern is not destabilized by adaptation, and remains as a steady state. We call this phase *stable attractor* phase.

**Figure 9:**
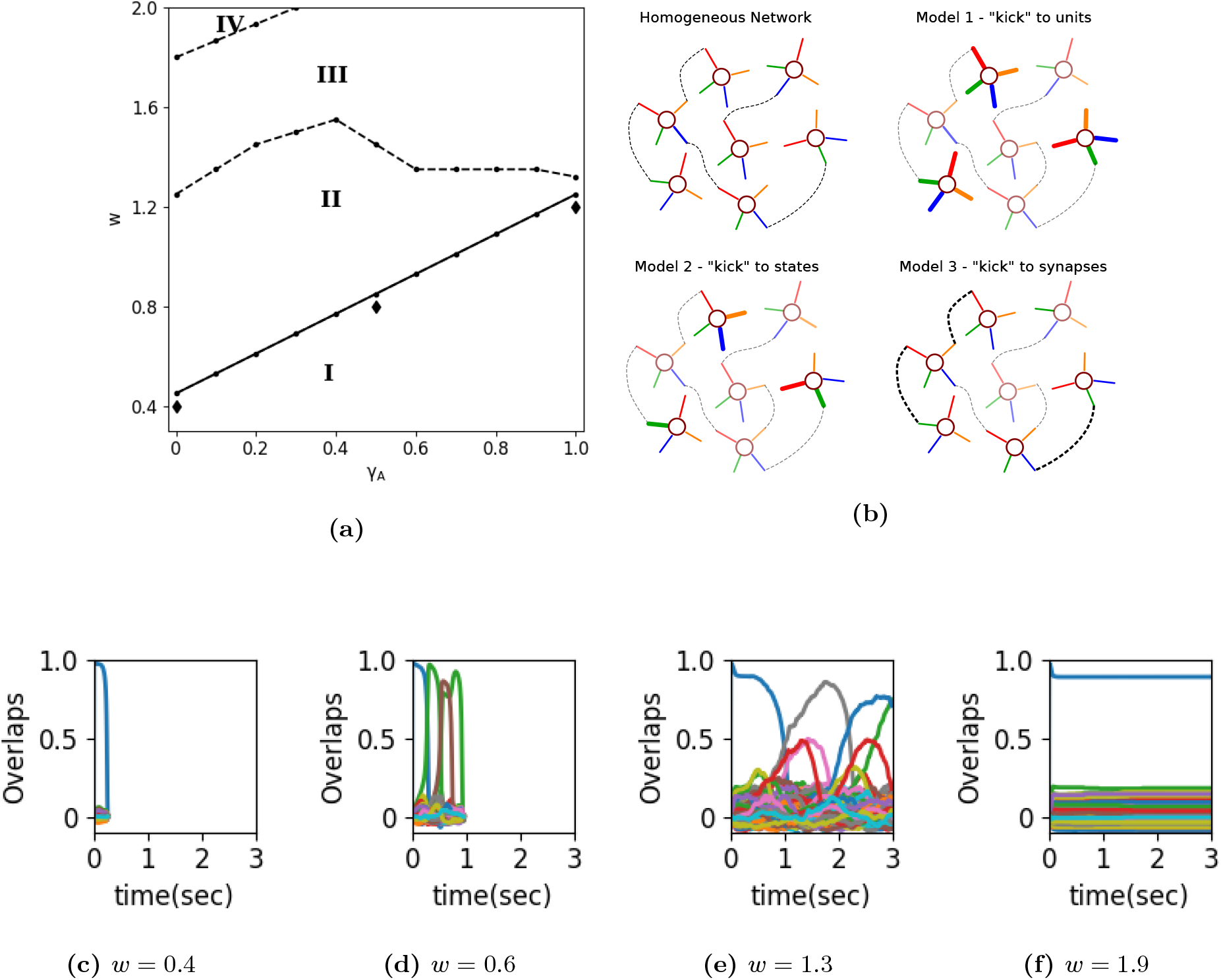
Moving on the phase diagram of the Potts model so it serves short-term memory. (a) Phase diagram of a Potts neural network in *w* − *γ_A_* plane. The *x*– axis is *γ_A_*, the proportion of fast inhibition. The *y* axis is the self-reinforcement parameter *w*. Increasing *w*, one observes different latching phases: no latching (I), finite latching (II), infinite latching (III) and stable attractor phase (IV). Diamonds indicate three points that are used in the study of short-term memory. **(c)** Distinct models of short-term memory are obtained by acting on different parameters, as explained in the main text. **(c)-(f)** Sample latching sequences with uncorrelated patterns of a network in the slow inhibition regime (*γ_A_* = 0, *τ*_1_ = 0.01*s*, *τ*_2_ = 0.2*s* and *τ*_3_ = 100*s*). The *x*-axis corresponds to time. The *y*-axis corresponds to the overlap, each colour for an item in long-term memory. **(c)** For too low *w*, in the no latching phase, there is only retrieval and the network cannot latch onto another pattern. **(d)** Increasing *w*, one reaches the finite latching phase, where the network retrieves a finite sequence of patterns, with high overlap. **(e)** Increasing *w* further, one reaches the infinite latching phase, where sequences are indefinitely long but where the quality of latching is degraded. The mean dwell time in a pattern is also increased compared with the finite latching regime. **(f)** Increasing *w* even further, one gets to the stable attractor phase, where the network retrieves the cued pattern and can not escape from that attractor. Network parameters are *N* = 1000, *S* = 7, *a* = 0.25, *c_m_* = 150, *U* = 0.1, *β* = 11, *p* = 200.

**Figure 10:**
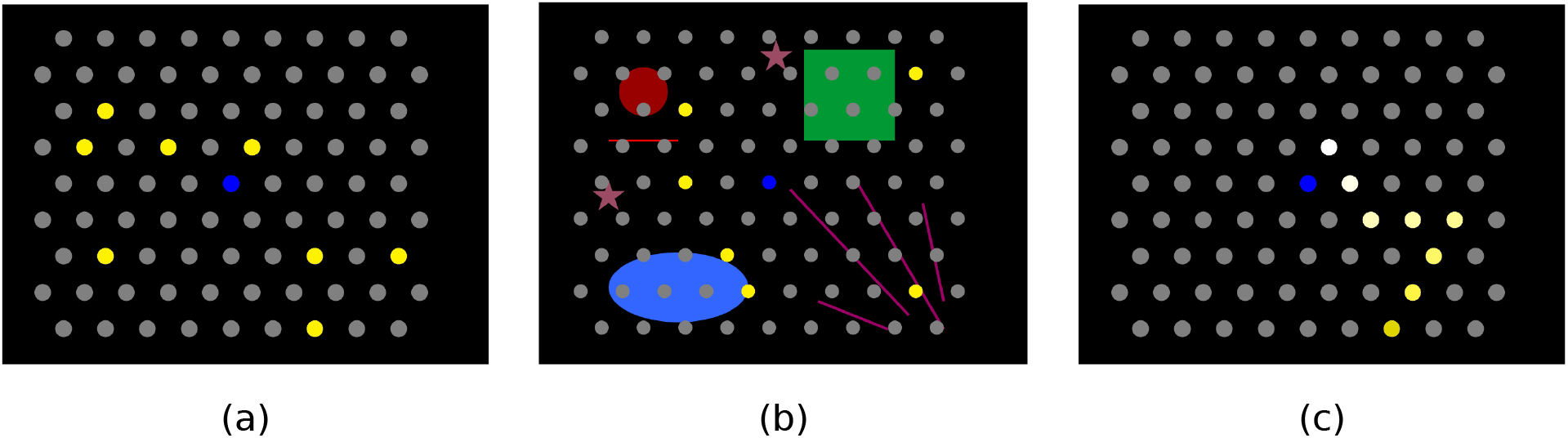
Sample stimuli used in the experiments. Participants were presented with a grid of grey dots on a screen, after which a series of yellow dots appeared on the screen. Subsequently after they had disappeared, they had to recall their locations by clicking on their positions. This experiment was carried out under several different conditions. **(a)** The dots appeared simultaneously and then disappeared all together. The participants then had to freely recall their positions **(b)** Same as in (a) but with additional landmarks, intended to probe whether landmarks help memory recall. **(c)** In this case, the dots appeared one by one (white to yellow) and formed a continuous trajectory, contrary to (a) and (b), after which participants performed serial recall.

The average dwell time in a retrieved pattern is continuously increasing with growing *w* in the infinite latching phase until the network enters the stable attractor phase. This is observed, though, only for *γ_A_* < 0.4, when other parameters are set as in Table 1. Except for this property, no major qualitative difference appears among the 3 regimes: slow, fast and intermediate inhibition regime.

**Table 1:**
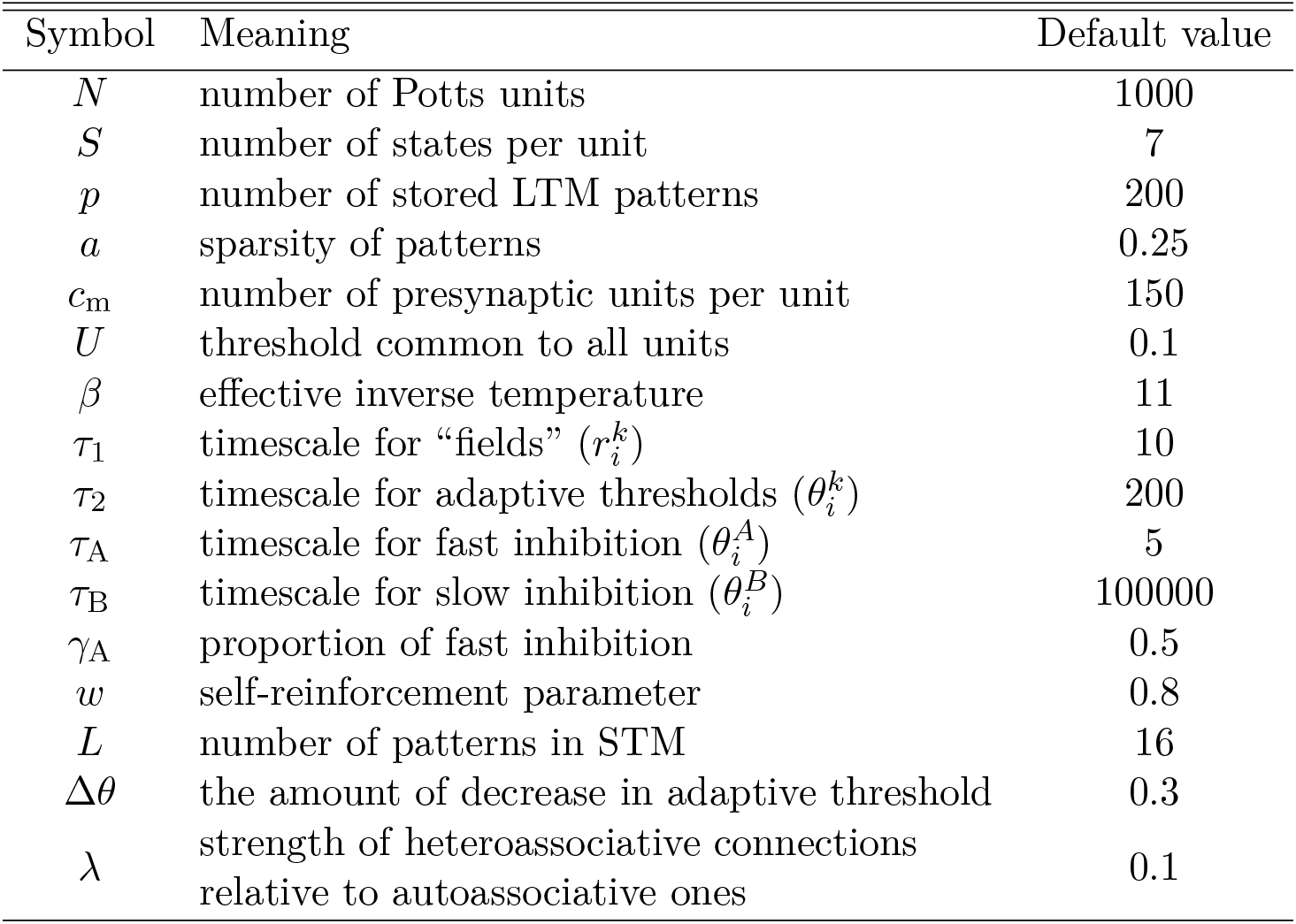
Parameters of the network

In [9], these phases and their phase transitions are studied in a semi-analytical way by using an *energy functional*, but only for the slow-inhibition regime. One can also use the same formalism to study phase transitions in the intermediate regime, where both slow and fast inhibition come into play. This study is still ongoing and will be reported elsewhere.

### 7.2 The Potts network as a model for short-term recall

The Potts network has been studied so far as a model of long-term memory; but it can be tweaked in minimal ways to serve also as a model of short-term or working memory. While it remains a simple or simple-minded object to study, it demonstrates how memory operating on widely different time scales can utilize the very same neural representations and the same associative mechanisms, based on plausible and *unsupervised* synaptic plasticity rules.

The core idea is that a few memory items, or sequences of items, are strengthened by increasing by a moderate and imprecisely determined amount the value of some pre-existing parameter, to effectively bring the network across a phase transition, into a phase in which those items or those sequences are held effectively separate from the ocean of all items and all possible sequences in long-term memory. So it is just an extra kick, without adding new components. The increase or extra kick is assumed to be temporary, and once it subsides, the short-memory has vanished. A critical assumption is that, since whatever plasticity in the brain serves as the extra kick, it has a transient time course, we model it by modifying parameters in simple and coarse ways, in contrast with what we assume to happen when encoding long-term memories, which in principle can be refined over many repetitions/recall instances, and can be taken therefore to reflect very precisely set parameters, down to the level of individual synaptic efficacies. Evidences and arguments supporting the model of short-term memory as an activated portion of long-term memory can be found in [1].

When a subject is performing a task of immediate recall, we posit the following assumptions.

- At the encoding phase, the attractors corresponding to the presented items are facilitated. We can visualize them as becoming wider and deeper in their basins.
- At the recall phase, we interpret that an item is recalled if the corresponding attractor is visited by the network.

The facilitation of attractors for STM items can be done by changing some parameters of the network. We propose in the main text three different models for short-term memory function that can constrain latching dynamics to a small subset of activity patterns that represent items in long-term memory.

### 7.3 The Potts model for serial recall

We use Model 2 to approximately constrain the dynamics to a subset of *L*_0_ patterns, for example the 6 digits of our experiment. We have *p* = 200 patterns in long-term memory, among which we give a Δ*θ* kick *L*_0_ = 6 patterns, indicated as 1, 2,…, 5, 6. In addition to the autoassociative connections between Potts units given by Eq. (6), we introduce heteroassociative connections to mimic the sequential order of the items presented in the experiment; we randomly pick *L* items among the 6 items (1, 2, 3, 4, 5, 6), allowing repetitions. When *L* = 6, for example, it can be 2 → 4 → 3 → 2 → 5 → 1. But we do not include sequences that have a subsequence like *AA* or *ABA* because the Potts model cannot really express such sequences (they occasionally appear in the dynamics, but only when the transition from A to B is incomplete or anomalous). We call sequences without any subsequence of the form ABA and AA *Potts-compatible*. In this way we prepare a set of 80 Potts-compatible sequences for a given value of *L*, with *L* = 3, 4, …, 10. If we denote a sequence of this set as *I*_1_, *I*_2_, …, *I_L_*, then the model for serial recall is determined by the following equations

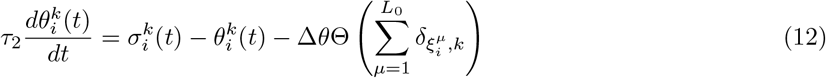

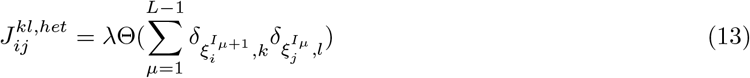

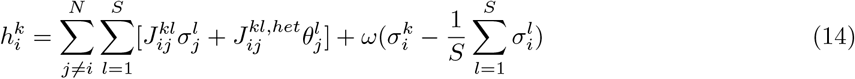

#### Quality of latching and correlation between patterns

The quality of latching is evaluated by means of *d*_12_ − *Q*. *d*_12_ is the difference between the largest overlap and the next largest one, averaged over time and over so called *quenched* variables [18], while

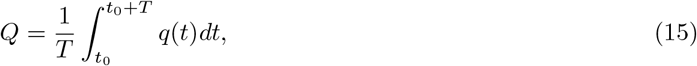

where 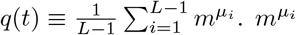 is the overlap of the network activity with a pattern *μ_i_* and *μ_1_*, …, *μ_L−1_* are the *L* − 1 patterns having largest overlaps excluding the maximum overlap. This quantity is a kind of measure on how “condensed”, i.e., partially recalled, the non-recalled patterns are.

The correlation between patterns is measured by two quantities [9, 33],

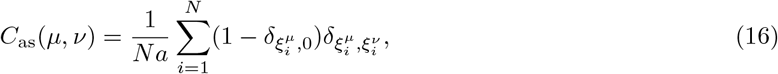

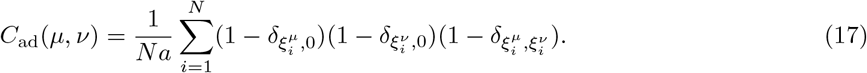

The average values of *C*_as_ and *C*_ad_ over different realizations of randomly-correlated patterns are given by

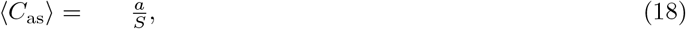

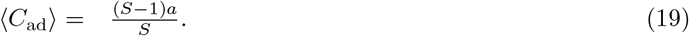

#### Network parameters

Network parameters used in this study are set as in Table 1, if not specified explicitly.

### 7.4 Experiments of free recall and serial recall

Both of the experiments were conducted online, with participants recruited through https://www.prolific.co/.

#### Serial recall

The 36 participants were instructed to watch a sequence appear on the computer screen and repeat the sequence just after, by clicking on the screen. They have to repeat sequences of *L* stimuli (*L* starting from 3). In one of the conditions, 5 trials for each length *L*, with *L* growing until 3 out of 5 trials are incorrect; the last *L* is then called the limit capacity for this participant in this condition. For each participant the sequences were of all three stimulus variants: - (D) Digits out of {1, 2, 3, 4, 5, 6} on a black screen, presented one at a time - (L) Locations on a hexagonal grid (Fig. 10) highlighted one by one, out of 6 around the central (blue) dot - (T) Trajectories on the same hexagonal grid: now each consecutively highlighted dot is one of 6 neighbors of the previous one (the first one is always one of the six around the center) Each stimulus was presented for one of the durations (in separate blocks): 400ms, 200ms, 100 ms. First always comes the 400 ms training session, then either 200 ms or 100 ms (balanced), and then the remaining duration. Presentation order was balanced across duration and stimulus material. In additional experiments, landmarks on the grid were used as well as intermediate presentation times, but no significant effect on the recall performance was observed.

#### Free recall

The same hexagonal grid as in serial recall is used (Fig. 10). In this experiment, the sets of stimuli were presented all at once, and the participants (*N* = 40) were instructed to repeat as many as they could recall, by clicking on the dots in the grid. For each set size *L* in {4, 6, 8, 12, 16, 24, 32}, the participants had 5 trials to do, each trial allowing for 2[*L* - (number of correctly recalled items)] clicks. For example, if participants correctly clicked 3 dots in a trial with *L* = 4, they were given another 2(4 − 3) = 2 clicks. Instead if they clicked incorrectly once and then correctly once, they had 2(4 − 1) − 1 = 5 clicks left. A set of size *L* was presented for 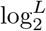 seconds.

## 8 Appendix Deriving scaling law under the assumption of equal visits

The quantity *M* can be estimated under the assumption of equal visits to each of the patterns. Under such an assumption, the probability of going to a new item *m* times and to one already visited at the *m* + 1-th time step is given by 1(1 − 1/(*L* − 1))…(1 − (*m* − 1)/(*L* − 1))*m/*(*L* − 1) and this contributes to *M* = *m* + 1. So taking a sum for *m* from 1 to *L* − 1 of this probability times *m* + 1 gives

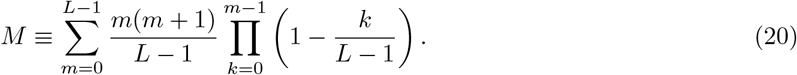

One simple approximation of this expression for *L* large yields

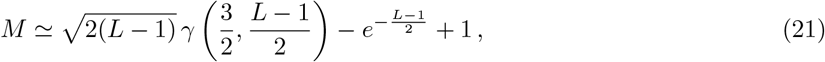

where *γ* is the *lower* incomplete Gamma function, which for *L* → ∞ grows as a square root,

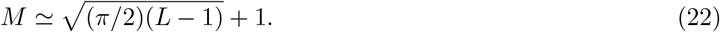

One way to approximate this expression for *L* large is to assume 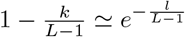, so that the product of the exponentials becomes the exponential of the sum, and one has

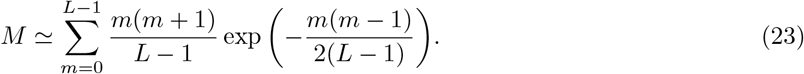

To further approximate the above sum with an integral, let 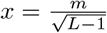, then we have

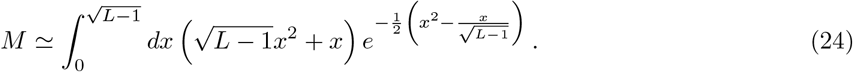

Keeping only the first term in the exponent of the integral

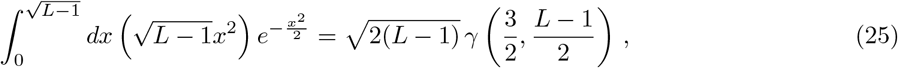

where

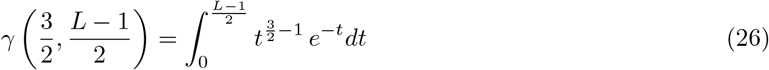

is the *lower* incomplete Gamma function, and

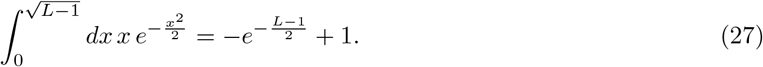

An alternative expression for *M* is in terms of the *upper* incomplete Gamma function,

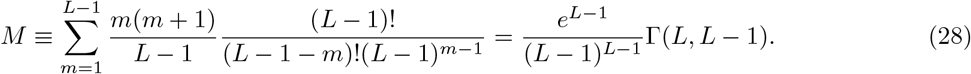

To derive its asymptotic behaviour for large *L*, it is convenient to separate one term and write

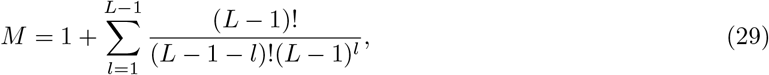

and then use Stirling’s approximation for the factorial to evaluate the sum as half an indefinite integral for −∞ < *l* < ∞, which can be evaluated at its saddle point near *l* = 1/2, yielding again, to leading order,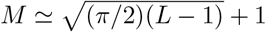

## Supporting information

Figs. S1 – S13 are in a separate file (PDF) named “Supplementary figures”.

## Acknowledgements

Work supported by the Human Frontier collaboration with the groups of Naama Friedmann and Remi Monasson on analog computations underlying language mechanisms, HFSP RGP0057/2016 and by EU Marie Skłodowska-Curie Training Network 765549 M-Gate.

## Author contributions

KIR conducted most of the analyses and simulations, which had been started by VB when still at SISSA, both with the supervision of AT, while OS run the experimental tests. All coauthors contributed to the evolving design of the study and to its write up.

## Conflict of interest

The authors declare no conflict of interest.

